# SimHumanity: Using SLiM 5.0 to run whole-genome simulations of human evolution

**DOI:** 10.1101/2025.09.01.673541

**Authors:** B.C. Haller, C.W. Nelson, M.F. Rodrigues, P.W. Messer

## Abstract

The reconstruction of human evolutionary history has undergone repeated advances, each made possible by methodological innovations. In recent decades, genetic and genomic data played a central role in the reconstruction of major evolutionary events such as the out-of-Africa migration, and genetic simulations of human evolutionary history have come to play a major role in testing more specific hypotheses including proposed patterns of migration and admixture with archaic hominins. Increasing computational power has allowed human evolutionary history to be modeled at ever-larger scales, but simulations that encompass the complete human genome, including sex chromosomes and mitochondrial DNA, have been difficult due to the lack of support for whole-genome models in commonly used evolutionary simulation frameworks. With the recent introduction of SLiM 5 such simulations are now straightforward to construct, allowing the easy simulation of humans at whole-genome scale under different demographic models and evolutionary dynamics. We here present three versions of a reusable, customizable, open-source SLiM 5 model for simulating the molecular evolution of the full human genome. We also show some simple analyses of results from the model, to illustrate its utility. We hope this model, which we have nicknamed “SimHumanity” in jest, will facilitate further progress in the field of human evolutionary simulations.

## Introduction

Modern progress in the study of human origins has been driven by a series of methodological innovations. Most notably, the advent of DNA sequencing allowed direct interrogation of the heritable genomic record that is present in every living human. Compared to other types of data, DNA data are relatively complete, objective, and relevant: for example, while fossils did not necessarily leave descendants, the genomes of all extant organisms did necessarily have ancestors [1].

As DNA sequencing technology advanced, genetic information could be queried at increasingly fine granularity, often overturning accepted wisdom. In the 1980s, morphological evidence had suggested that humans are most closely related to gorillas [2]; DNA evidence showed that humans are more closely related to chimpanzees [3,4]. Similarly, paleontological evidence had supported a ‘multiregional model’ in which isolated human populations evolved in parallel [5]; DNA evidence showed that all humans share a recent origin in Africa [6,7]. Later, the first fragmentary DNA evidence from Neanderthals had suggested they were simply replaced by *Homo sapiens*; genomic data instead showed they interbred with non-African humans [8].

Subsequent research has led to the discovery of Denisovans [9] and established the likelihood not only of archaic hominin introgression and admixture, but also genetic exchanges between ancient human subpopulations in Africa [10] and ancestry contributions from so-called “ghost” populations [11–13].

As evolutionary and demographic hypotheses about human history have increased in complexity, the toolbox of methods for evaluating these hypotheses has come to require simulation. The earliest simulations involved a backward-in-time approach, where a set of sequences is traced to their ancestors until a generation is reached in which all lineages converge to a single common ancestor. This so-called ‘coalescent’ framework therefore only models ancestors of a sample – not the many more individuals who left no descendants in the given sample. Coalescent simulations are computationally efficient and have proved useful for many problems, including inference of human demographic history based on site frequency spectra with methods such as ∂a∂i [14], fastsimcoal [15], and fastsimcoal2 [16]; inference based on patterns of linkage with methods such as PSMC [17] and MSMC [18]; and inference based on both of those together as in SMC++ [19]. At the same time, the nature of the coalescent approach means that it is limited to simplistic evolutionary scenarios. We are now at a stage where insufficient model complexity, not insufficient data, is the main limit to progress.

Forward-in-time individual-based simulation, by contrast, involves explicit modeling of sequences starting from an ancestral population and following their descendants. Thus, this approach models all organisms alive in each generation – not limited to the unknown subset whose lines of descent will persist to the final generation – and relies upon fewer assumptions than the coalescent. Such simulations can implement arbitrarily complex scenarios involving any number of parameters governing aspects of the simulation such as mutation, selection, demography, genetic architecture, linkage, and geographic spatiality (among others), which can be tested in unlimited combinations. Results are then compared with empirical data, and the degree of agreement between empirical and simulated results is used to assess the plausibility of the model.

The forward-in-time approach, while flexible, has until recently been computationally prohibitive for whole-genome scenarios in reasonably large populations; explicitly simulating every mutation in every individual is inevitably costly in terms of both computation time and memory usage. But with advances in computational power, from the onward march of Moore’s Law to the increasing availability of computing clusters, modeling whole genomes has become increasingly feasible depending upon factors such as population size, genome size, and the number of generations to be modeled. In response to this trend, some software models for running individual-based simulations have expanded their scope, extending their capabilities to take advantage of the greater computational power available.

Here we will focus on one software package that exemplifies this trend: the evolutionary simulator SLiM, which some of us (Messer and Haller) have developed. The original SLiM 1 [20] focused on simple Wright–Fisher models of a single chromosome, configured by a structured input file that expressed simple events such as demographic changes. SLiM 2 [21] added scriptability, providing much greater flexibility, and provided a new graphical modeling environment called SLiMgui for interactive model development and exploration. Furthermore, SLiM 2.3 added support for spatial models, including spatial interactions between individuals and their neighbors. In SLiM 3 [22] the internal model was extended to support non-Wright– Fisher models, allowing much greater individual-level detail and biological realism, as well as support for “tree-sequence recording” [23], the recording of genetic ancestry trees along the chromosome. Furthermore, SLiM 3.3 added support for explicit modeling of nucleotide sequences (as opposed to SLiM’s default, a more abstract representation of mutations without nucleotides). SLiM 4 [24] added support for multiple species, and ecological interactions between species, broadening SLiM’s ambit from population-genetics simulations to include evolutionary ecology simulations as well. In all of these advances, the common thread is that SLiM’s capabilities were extended and expanded to take advantage of the increased computational power newly available to many researchers even on a single laptop, making it possible to build larger and more realistic simulations of evolutionary and ecological dynamics.

Earlier versions of SLiM have already proved useful in studies of human evolution. Many of these studies implemented the Gravel et al. (2011) model [25] of human demographic history [26–30], including the version of that model provided in SLiM recipe 5.4 [31]. However, due primarily to computational limitations, much of this work was limited to a single representative autosomal genomic region per individual, with no modelling of sex chromosomes or mitochondrial DNA. Two notable exceptions are Rinker et al. (2020) [28], who modeled the full autosomal genome but collapsed the intergenic regions to single sites with recombination rates proportional to their original length; and Santander and Moltke (2025) [31], who modelled the full lengths of the two shortest chromosomes, 21 and 22, including intergenic regions, for a total of ∼68.9 Mbp per individual.

In these older versions of SLiM, modeling multiple chromosomes was not intrinsically supported. Both Rinker et al. (2020) [28] and Santander and Moltke (2025) [31] worked around that limitation by using a recombination rate of 0.5 at specific points in the simulated genome, producing “effective chromosomes” that were unlinked from each other, although SLiM did not actually “understand” that separate chromosomes were being modeled. However, this technique only works to simulate multiple autosomes; for other types of chromosomes, such as sex chromosomes, substantial technical scripting would be necessary to manage the inheritance and recombination of the non-autosomal chromosomes. The complexity of that scripting made writing whole-genome models in older versions of SLiM very difficult; to our knowledge, no published studies have done this.

The most recent major release, SLiM 5 [32], fundamentally improves upon SLiM’s ability to model whole genomes by providing intrinsic support for models that include multiple autosomes, sex chromosomes, and even other genomic entities such as mitochondrial and chloroplast DNA, making it trivial to do what was previously almost prohibitively difficult. All of the details of inheritance, including recombination, are handled by SLiM automatically in such models. Multi-chromosome models are more computationally intensive than single-chromosome models, of course, since a much larger amount of genetic information is simulated; but it is nevertheless possible to construct whole-genome models of human evolutionary history that can run in a reasonable amount of time on typical hardware such as a laptop.

Modelling entire genomes is important because evolutionary processes shape variation at broader scales. Recombination rates vary considerably within and between chromosomes [33], as does the density of targets of selection (e.g., coding sequences) [34]. Mutation rates also probably vary at this scale [35]. Nevertheless, population genetic simulation studies have usually focused on modelling a few hundred kilobases to megabases of sequence. Only recently have we begun to appreciate the need for modelling at larger scales. For example, Gower et al. (2025) [36] demonstrated that the power to detect selective sweeps is highly heterogeneous along a chromosome. The fact that evolutionary processes are heterogeneous along genomes can also be a blessing: Rodrigues et al. (2024) [37] showed that large-scale patterns of genetic variation in the great apes contain information about the relative roles of background selection, positive selection, and mutation rate variation.

To facilitate whole-genome simulations of humans in SLiM, we have developed a reuseable, open-source SLiM model named “SimHumanity” in jest. All models are simplifications, and this one is no exception; nevertheless SimHumanity serves as a common starting point for contexts in which simulating the whole genome is useful. We will show three different versions of our model, to demonstrate how it can be customized for different purposes. We will briefly present results from each of these model variants, to illustrate their behavior and utility for pedagogical purposes. In two of the three versions of our SimHumanity model we implement the aforementioned Gravel model of human evolutionary history [25], which was developed by applying ∂a∂i [14] to early 1000 Genomes data [38]. The Gravel model has been extended in myriad ways, such as the inclusion of African and additional European population expansion [39] and more detailed Asian population substructure [40]. Despite its simplicity, the Gravel model still serves as a common and useful baseline for modeling human history, and can be modified to suit the investigator’s particular research question.

### Model 1: the foundation

The first version of the SimHumanity model that we present (Code Sample 1) simulates both neutral and non-neutral mutations on 25 chromosomes: 22 autosomes, an X chromosome, a Y chromosome, and the mitochondrial genome. (Note that when separate sexes are modeled, certain chromosomes in certain individuals – like the Y in females – are represented by null “placeholders” to accurately reflect karyotypic sex differences; see below.) It simulates a trivial demographic model consisting of a single subpopulation of *N*=1000 individuals for 10*N* generations. This version of the model serves primarily as a demonstration of how to set up the genetic structure of a multi-chromosome model in SLiM 5; we will walk through it here step by step.

The genetic configuration of the SimHumanity model – chromosome lengths, recombination rate maps, genome annotations providing coordinates of coding sequences, the distribution of fitness effects (DFE), and so forth – comes from a variety of sources. The sources of these data and the ways in which they were collected are detailed in the “Genetic structure data” section below; here we will take that data as given, and focus on the construction of the model itself. All versions of the SimHumanity model presented here use the same genetic configuration, except for specific modifications discussed.

The model begins with an `initialize()` callback, a block of code that runs once at the beginning of the simulation to initialize the model’s configuration. It begins by calling `defineRepositoryPaths()`, a user-customizable function that finds the data repository for this paper at a given path, and checks that particular sub-paths exist within that repository. The code for this function is provided in the repository version of this model, but is omitted here for brevity.

Then the simulation configuration begins with a call to `initializeSLiMOptions()` to tell SLiM that this is a nucleotide-based model; in consequence, every mutation will have an associated nucleotide, and every chromosome will need an ancestral sequence that provides the reference nucleotide in positions where a mutation is not present. A call to `initializeSex()` enables separate sexes in the model (otherwise the default would be hermaphrodites, and sex chromosomes would not be allowed), and a constant `N` is defined as `1000`, representing the number of individuals that will be simulated. Several more constants are defined to represent the mutation rates associated with the `PosNeg_R24` DFE, as discussed in the “Genetic structure data” section; `MU_TOTAL` is the overall mutation rate, `MU_BENEF` and `MU_DELET` are the beneficial and deleterious rates, and `MU_NEUTR` is the neutral mutation rate within coding regions – the rate that remains after the beneficial and deleterious rates are subtracted from the overall mutation rate.

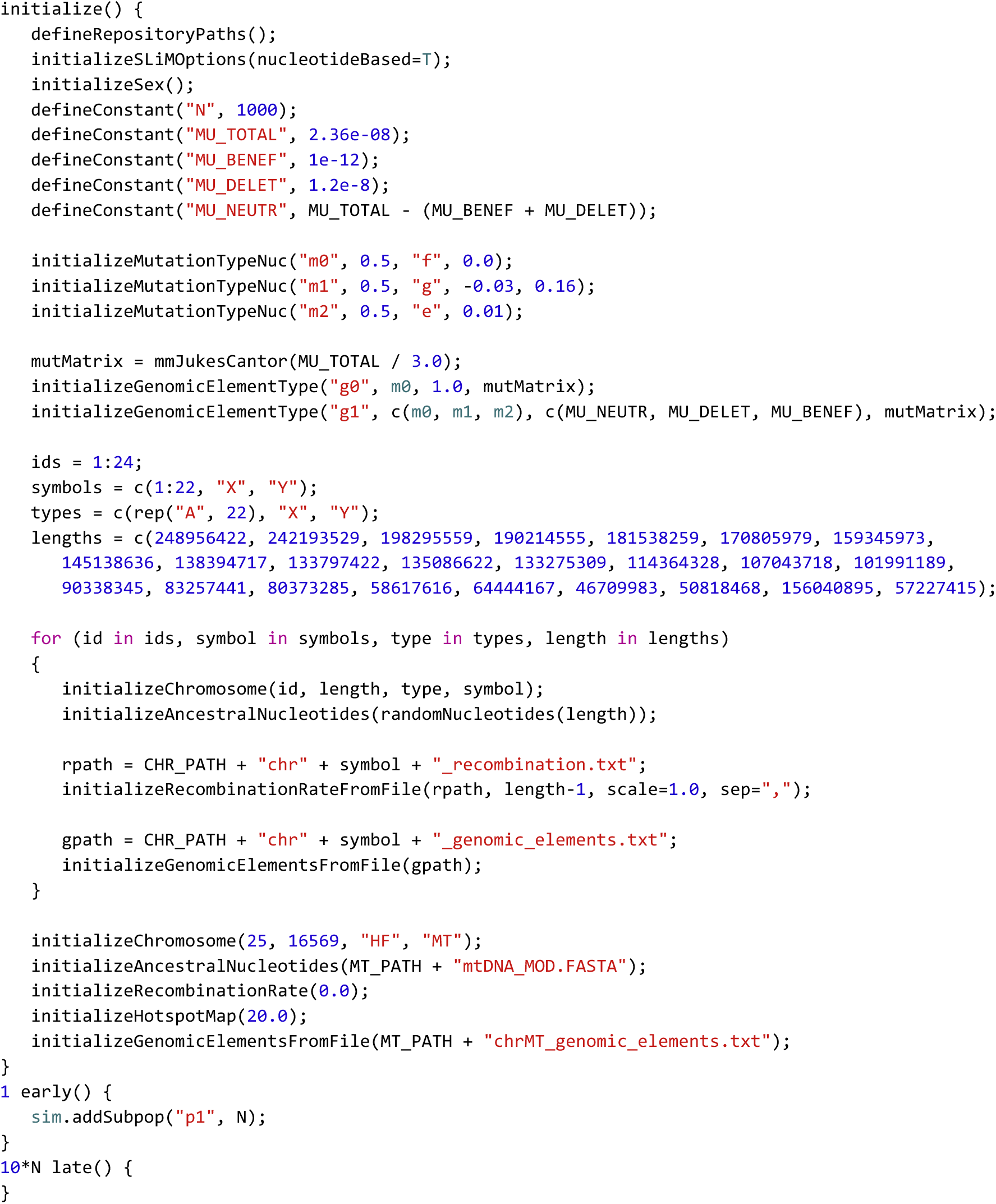

**Code Sample 1.** The foundational SimHumanity model, containing both neutral and non-neutral mutations simulated for all human chromosomes (including mtDNA) in a population of size *N*=1000, for 10*N* (10,000) generations. This model is provided in our repository in the file `model1_foundation.slim`; that file also contains comments in the code, which have been removed here for brevity. Instead of comments here, this code is discussed in the text. The functions `defineRepositoryPaths()` and `initializeGenomicElementsFromFile()` are also omitted here, but their definitions are present in the version of this file in the repository.

Next we have three calls to `initializeMutationTypeNuc()` to set up “mutation types” that represent DFEs from which neutral (`m0`), deleterious (`m1`), and beneficial (`m2`) mutations will be drawn. The first mutation type, being neutral, draws from a fixed (type `"f"`) DFE with a selection coefficient of `0.0` (and a dominance coefficient of `0.5`, which does not matter for neutral mutations). The second draws from a gamma distribution (`"g"`) with a specified mean and shape, and the third draws from an exponential distribution (`"e"`) with a specified mean; these both also use a dominance coefficient of `0.5` (semidominant).

The next three lines set up “genomic element types” representing types of regions within the genome; in this model we define just two genomic element types, non-coding (`g0`) and coding (`g1`). Non-coding regions draw their mutations only from `m0`, the neutral mutation type; coding regions draw their mutations from `m0`, `m1`, and `m2`, in proportions specified by the `PosNeg_R24` DFE as discussed previously. Both genomic element types use a mutation rate matrix provided by the Jukes–Cantor model; mutations therefore occur at a constant rate regardless of sequence, with equal probability for each possible nucleotide at a new mutation. The *α* parameter passed to `mmJukesCantor()` is used by the Jukes–Cantor model in such a way that the overall mutation rate is 3*α*; given that the required input is the value of *α*, we therefore supply one-third of the desired total mutation rate to this function.

With that overall structure configured, SLiM now knows how to model both non-coding and coding regions. Most of the rest of the code configures chromosomes to be sequences of such regions. A joint `for` loop iterates over the chromosome ids (integer identifiers), symbols (string identifiers), types (such as `"A"` for autosome, `"X"`, and `"Y"`), and lengths (in base positions), in synchrony. Each iteration of the loop therefore configures one chromosome, based upon the values of the four loop index variables. First, a call to `initializeChromosome()` provides SLiM with that basic information – its `id`, `symbol`, `type`, and `length`. This creates the new chromosome object and prepares it for further configuration. Next, a random ancestral nucleotide sequence is generated and set; given size considerations, the exact sequences are not provided here, but these could be downloaded from NCBI and loaded into SLiM by providing the FASTA file paths to `initializeAncestralNucleotides()`, as demonstrated for the mitochondrial genome below. Then the recombination rate map for the chromosome is loaded from a file, and the “genomic elements” that define particular non-coding and coding regions along the chromosome are loaded from another file; these files are provided in our repository as described in the “Genetic structure data” section. In this manner, each chromosome is configured in turn. Note that the `initializeGenomicElementsFromFile()` function is not built into SLiM; it is a user-defined function, defined as a part of this model’s script but not shown here for brevity (it is provided in the repository version of this model).

The last five lines of the `initialize()` callback similarly initialize the haploid chromosome that represents the mtDNA. The steps here are essentially the same as for the other chromosomes; they are just done separately here because the mtDNA structure data is located in a different place in the repository, as discussed in the “Genetic structure data” section. For the mitochondrial DNA we do load the ancestral sequence from a FASTA file, demonstrating that technique. The chromosome type `"HF"` stands for “haploid female”, and means that the mitochondrial DNA is haploid and will be inherited clonally from the female parent, but will be present in both sexes; SLiM provides twelve different chromosome types, in fact, but we will focus on `"A"`, `"X"`, `"Y"`, and `"HF"` here since they are the types we need to model humans.

There is one twist here, however: mitochondrial DNA typically has a higher mutation rate than autosomal DNA, because there is less error-correction in the copying process; as discussed in the “Genetic structure data” section, we use a rate of 4.0e-7 mutations per base position per generation in this model, averaged from the results of two recent papers. Our model uses a Jukes–Cantor mutation rate matrix as explained above, but in nucleotide-based models in SLiM, the mutation rate can also be sequence-dependent to allow, for example, elevated mutation rates at CpG sites. Further, to express mutational hot spots and cold spots in the genome, one can define a “hotspot map” that multiples the sequence-dependent rate by different scaling factors in different regions, scaling the mutation rate up or down as desired. To account for the elevated mtDNA mutation rate, we have simply increased the mutation rate in the mtDNA chromosome using a hotspot map, essentially just designating the entire mtDNA chromosome as a “hot spot” (equivalently, we could have defined another Jukes–Cantor mutation rate matrix with a different rate). If we had mutation rate map data for the chromosomes being modeled, indicating hot and cold spots, we could similarly use those data to define hotspot maps for each chromosome, incorporating fine-grained mutation rate data into the model. For now, however, we just use the hotspot map to elevate the mutation rate for the entire mtDNA chromosome using a multiplier of 4.0e-7 / 2.0e-8 = 20 times the baseline mutation rate used in the other chromosomes. This is the purpose of the `initializeHotspotMap()` call; it scales the mutation rate for the mitochondrial chromosome up by a factor of 20 relative to the baseline rate defined by the Jukes–Cantor mutation rate matrix.

After that, an event for SLiM to execute is defined with `1 early()`, meaning that this event will be run early in generation 1. This event adds a new subpopulation called `p1` with `N` individuals. This event therefore triggers the execution of the SLiM simulation by creating a population of individuals that will evolve, according to the Wright–Fisher model, with mating, births, deaths, mutations, recombinations, and so forth. And finally, a `10*N late()` event – thus defined as running late in generation `10*N` – defines the endpoint of the simulation, since it is the last event scheduled in the model. This is where one would put calls to produce final output, but here we simply allow the model to end. The final state of the model can easily be observed visually in SLiMgui, SLiM’s interactive modeling environment.

Since this model includes every mutation on every chromosome, it generates quite a large amount of genetic information, and therefore takes a fairly long time to run; on a test machine (a Mac Studio 2022, M1 Ultra, 128 GB memory) a run of the model at the command line took 7448 seconds (2.07 hours) with a peak memory usage of 7.47 GB. The performance of this model, in terms of both runtime and memory usage, is perhaps acceptable at its population size of 1000, and it is fast enough that a set of replicate runs could easily be conducted on a computing cluster; but it would have trouble scaling up to very large population sizes, and it would be hard to explore the behavior of this model comprehensively across a range of parameter values such as multiple mutation rates or putative population histories. Better performance is therefore desirable, and we will see techniques for achieving that in the following sections.

This model runs for 10*N* generations because the intention is for it to execute a burn-in period: a period of simulation that builds up to an equilibrium level of genetic diversity. (Alternatively, the burn-in period can be viewed as building relatedness between the simulated individuals until they reach coalescence.) For a simple single-subpopulation neutral Wright–Fisher model, an often-quoted rule of thumb called the “10*N* rule” says that the burn-in should be at least 10*N* generations to reach something close to mutation–drift equilibrium. To illustrate this, we added extra code to this model (provided in the repository as `model1_foundation_FIGURE.slim`) that uses SLiMgui’s built-in custom plotting functionality to produce site-frequency spectrum (SFS) plots for the model over time (Fig. 1). Early in the run, there is an excess of mutations at low frequency and a deficit at high frequency, because the model starts out with no genetic variation at all; all of the variation present early on comes from new mutations that begin as singletons and have not yet had much chance to rise in frequency. As the model runs, the observed SFS converges towards the expected SFS.

**Figure 1.**
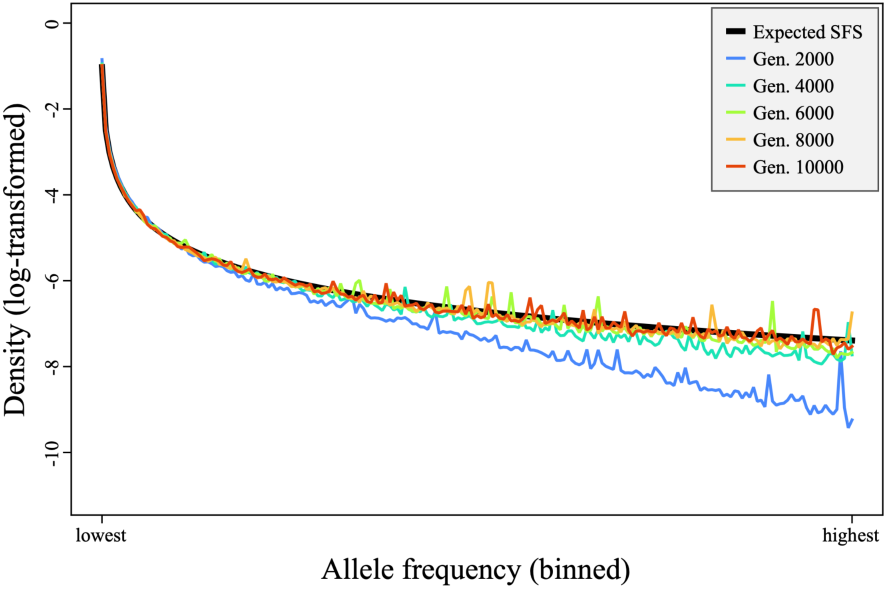
Expected versus observed site frequency spectra (SFS) over time. The thick black line is the expected SFS, and the thinner colored lines show the observed SFS at different time points (see legend). The model begins with a deficit of high-frequency derived alleles, and converges towards the expected SFS over time. The unfolded SFS is summarized into bins containing 10 count values each (to decrease noise), ranging from the ten lowest frequencies (counts of 1 to 10) to the ten highest frequencies plotted (counts of 2*N*−19 to 2*N*−10, where *N* is the population size); the nine highest counts, from 2*N*−9 to 2*N*−1, are not plotted since that bin would contain only nine values. The binned SFS is then normalized to densities summing to 1.0, and then log-transformed (to visually emphasize bins with low density). This plot was generated by SLiMgui running `model1_foundation_FIGURE.slim`, a modified version of model 1 provided in our repository.

It’s worth noting, however, that this model will not converge *exactly* upon the expected SFS, because it is not a neutral model; we have (common) deleterious mutations and (rare) beneficial mutations in this model, both of which will influence the SFS. Nevertheless, Fig. 1 illustrates that the model run begins out of equilibrium, and that a burn-in period is needed for its genetics to reach an equilibrium state before other phenomena can be appropriately modeled. The necessary burn-in period for this non-neutral model would no longer be 10*N* – and indeed, 10*N* is only a rough rule of thumb even for simple neutral models, and should be taken with a grain of salt [31]. Using techniques such as plotting the SFS over time, as shown here, it should be possible to observe approximately when equilibrium has been reached for a particular model.

The run of model 1 shown in Fig. 1 was conducted in SLiMgui, which generated the plot shown. At the end of the run there were 1,912,662 mutations segregating (of which 99.57% were neutral) and 386,014 fixations (of which 99.74% were neutral). The high percentage of neutral mutations observed is unsurprising, since ∼99% of the genome is neutral non-coding regions, and since deleterious mutations are more likely to be lost than neutral mutations. The large number of neutral mutations means that SLiM spends quite a lot of time on internal mechanics, particularly recombination.

### Model 2: adding demography

The second version of the SimHumanity model that we present (Code Sample 2) simulates only non-neutral mutations, omitting neutral mutations. It again simulates all chromosomes, with the same genetic structure as model 1. The main biological change to the model is that it now implements the demographic model of Gravel et al. (2011) [25]. This version of the model serves primarily as a demonstration of how to set up a demographic model, and also shows that omitting neutral mutations can sometimes be useful for performance. We will examine its differences compared to model 1; much of its initialization code is the same.

The `initialize()` callback is modified so that neutral mutations are no longer simulated; mutation type `m0` and genomic element type `g0` are no longer used, and genomic element type `g1` now has a mutation rate, defined as a new constant `MU_CODING`, that reflects the lower total mutation rate due to the omission of neutral mutations. This change eliminates approximately 99.57% of segregating mutations (as seen with model 1), but there will still be a great many deleterious mutations in the model, and they take much more of SLiM’s time per mutation than neutral mutations because their fitness effects have to be evaluated.

The other change is the addition of the demographic model of Gravel et al. (2011) [25], which we will walk through now. Note that the SLiM manual [41] contains a more extensive discussion of this model; those who are interested in its specifics should refer to that resource and the original paper. Here we are interested in simply using it as an example of integrating demographic events into the SimHumanity model.

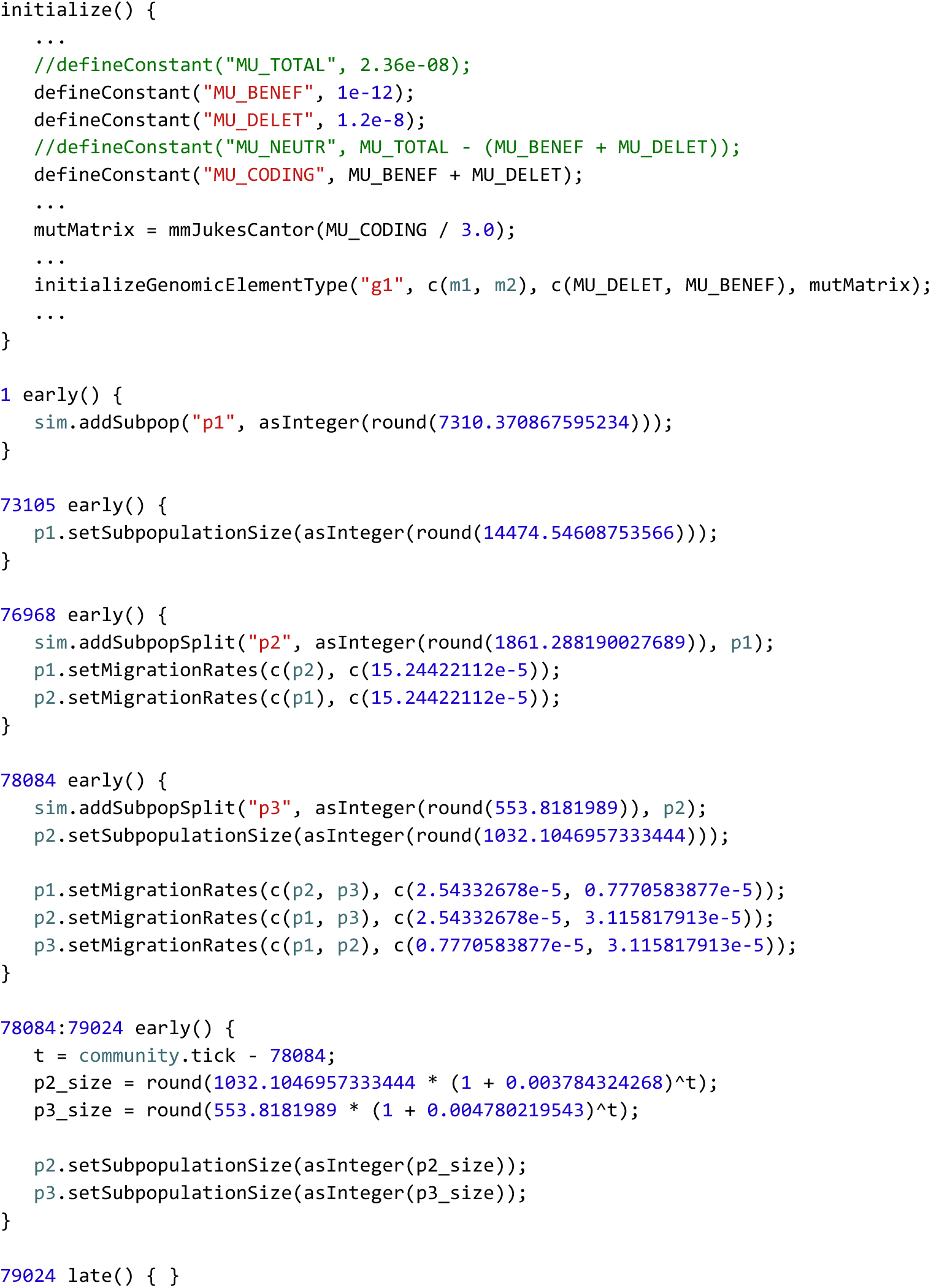

**Code Sample 2.** This version of the SimHumanity model contains only non-neutral mutations, again simulated in all chromosomes. It implements the Gravel et al. (2011) [25] demographic model, as discussed in the text; that code replaces the trivial demography code in model 1. Only differences from model 1 are shown here; the full model (again, with comments that are omitted here for brevity) is provided in `model2_demography.slim` in our repository. Mutation type `m0` (neutral) and genomic element type `g0` (non-coding) are no longer used, and the user-defined function `initializeGenomicElementsFromFile()` is modified from model 1 to only create `g1` (coding) genomic elements, but is not shown here (just as it was not shown in Code Sample 1). Double forward slashes (`//`) represent comments (non-executed code) in a SLiM model.

The model begins in `1 early()`, as before, by creating subpopulation `p1`. This represents the African population that is hypothesized to be ancestral to all of humanity. It is created with a size of `7310` individuals, which represents the effective population size (*N*_e_) estimated by Gravel et al. (2011) [25], not the population’s census size, but is used as the census size here as an approximation. Here and elsewhere in this model, population sizes are expressed as rounded, integer versions of the exact parameter estimates from the original model.

A series of `early()` events then steps through the demographic model. The first of these executes in generation `73105`. This time point marks the end of the model’s burn-in period (following the 10*N* rule of thumb, although in this non-neutral model that is likely inexact as discussed for model 1). At the end of burn-in, the African population expands to a size of `14475`, approximately doubling its *N*_e_, and then continues to evolve.

In generation `76968` a new subpopulation, `p2`, splits off from `p1` with a size of `1861`; this represents the migration of humanity into Eurasia. A symmetrical migration rate is set from `p2` into `p1`, and from `p1` into `p2`; in SLiM, this rate expresses the expected fraction of the children in the destination subpopulation that will subsequently be generated from parents in the source subpopulation.

The population proceeds in that fashion until generation `78084`, when the Eurasian (`p2`) subpopulation splits into European (still `p2`) and East Asian (`p3`) subpopulations. The new `p3` subpopulation is given a size of `554` individuals, and `p2` is adjusted to a size of `1032`. New migration rates are set that exchange migrants between all three subpopulations at various rates.

In the same generation, exponential growth begins in the `p2` and `p3` subpopulations, implemented in the `78084:79024 early()` event. That exponential growth continues until the present day, which is generation `79024` in the model. It is implemented simply by calculating the new subpopulation sizes in each generation, with exponential functions, and setting those sizes for the subpopulations.

A final late() event in generation 79024 marks the end of the model, although strictly it is unnecessary; the model would end in generation 79024 anyway, since there are no further events scheduled to occur after the end of the exponential growth period.

A run of this model at the command line took 69,739 seconds (19.37 hours), with a peak memory usage of 25.8 GB (on the same test machine). This is more than nine times the runtime of model 1; the population size and the number of generations are both more than seven times higher than in model 1, so the performance benefits resulting from omitting the neutral mutations have been more than counterbalanced by the increased scale of the model. Still, omitting the neutral mutations made a substantial difference; a run of this model that includes neutral mutations (given as `model2_demography_WITH_NEUTRAL.slim` in our repository) took 132,936 seconds (36.93 hours), almost twice as long, with a peak memory usage of 48.2 GB.

To illustrate the potential utility of this model, even though neutral mutations were omitted from it, we again constructed a version of the model (`model2_demography_FIGURE.slim` in our repository) with additional code to generate a plot at runtime (Fig. 2). This plot shows every beneficial mutation that fixed during the run of the model. (It would also be possible to plot beneficial mutations that were lost, but that is not shown here.) It can be seen that the selective sweeps of beneficial mutations destined for fixation in this model very often overlapped in time, including instances of ‘stochastic tunnelling’ in which later beneficial mutations arise on (or recombine into) haplotypes that already contain earlier beneficial mutations [42,43].

**Figure 2.**
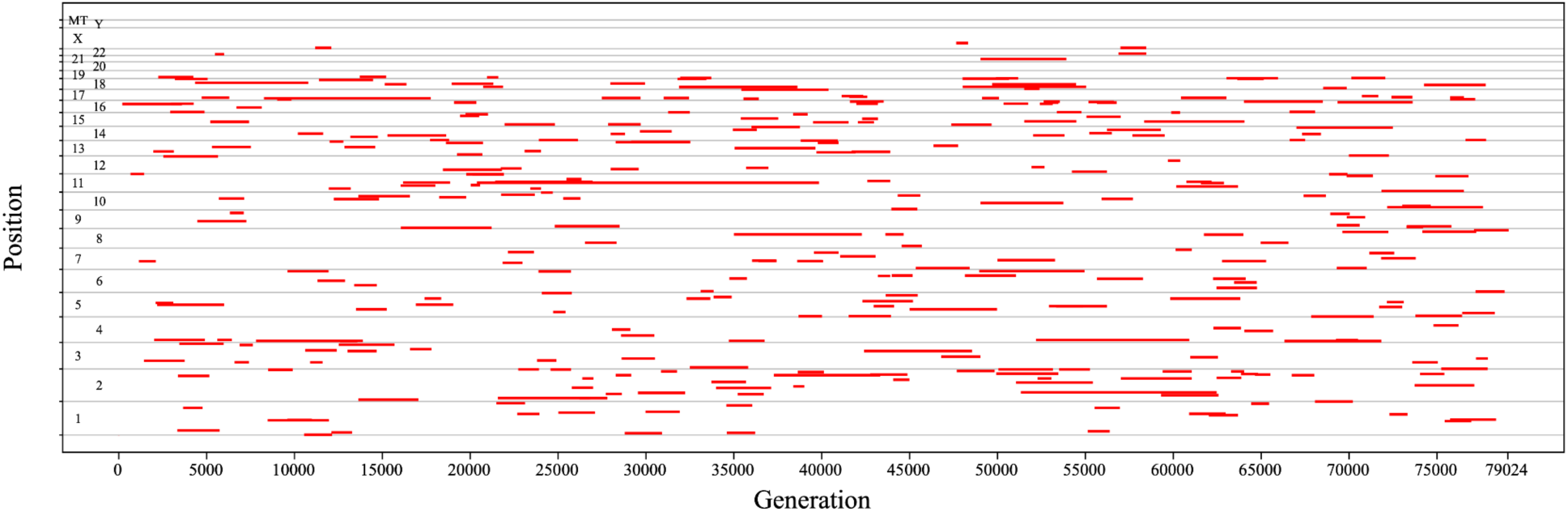
A history of all beneficial mutations that reached fixation during one run of model 2. The x axis represents time, in generations, and the y axis represents position in the genome (subdivided into the individual chromosomes simulated, which are labeled). Each red line represents a single beneficial mutation, spanning the interval from its generation of origin to its generation of fixation. It can be seen that the selective sweeps of mutations often overlap in time; indeed, it is common for several sweeps to occur simultaneously, even within a single chromosome, indicating that sweeps are probably affecting each other’s trajectories and probabilities of fixation. This plot was generated by SLiMgui running `model2_demography_FIGURE.slim`, a modified version of model 2 provided in our repository.

At the end of the SLiMgui run of model 2 shown in Fig. 2 there were 625,065 mutations segregating (99.98% deleterious, 0.0152% beneficial) and 6028 fixations (94.77% deleterious, 5.22% beneficial). The presence of so many deleterious mutations helps explain why the runtime of the model remains so long; evaluating their fitness effects in every individual in each generation is time-consuming. That overhead is largely not present for neutral mutations, since SLiM does not need to calculate their fitness effects, so the runtime overhead per deleterious mutation is much larger than the runtime overhead per neutral mutation.

One important thing to note is that the demographic parameters of the Gravel et al. (2011) [25] model were estimated by fitting the model to empirical data, and a purely neutral DFE was used to conduct that fitting. We have simply grafted that demographic model onto our genetic model, which contains beneficial and deleterious mutations; the demography would therefore no longer be expected to produce results that exactly fit the empirical data.

### Model 3: adding tree-sequence recording

Model 1 simulated neutral mutations as well as deleterious and beneficial mutations, which was quite slow. Model 2 did not simulate neutral mutations, but since the larger population sizes allowed so many deleterious mutations whose fitness evaluation is time-consuming, the performance was still not ideal as discussed previously. In this section we will add tree-sequence recording to the model, to see its benefits for the runtime performance and memory usage of the model.

Tree-sequence recording is a method for tracking the local ancestry tree at each position along a simulated chromosome, recording that information into a data structure called a “tree sequence” [23,44]. In a single-chromosome model, this produces a file on disk called a `.trees` file (because that is the filename extension conventionally used). In a multi-chromosome model, a directory containing one `.trees` file per chromosome is produced, called a “trees archive”. One can then work with these tree sequences in Python, either one chromosome at a time or jointly, using the `tskit`, `msprime`, and `pyslim` packages, as we will see.

Tree-sequence recording entails extra work to record the necessary information as the simulation runs. In addition to that recording, there is also substantial additional work in the form of a process called “simplification” that has to be done periodically to discard recorded information that corresponds to extinct branches of the evolutionary tree and other such information that is irrelevant to the final result; without periodic simplification, the size of the tree sequence often grows too large for practical purposes. However, the additional overhead of tree-sequence recording is typically more than compensated for by two optimizations that tree-sequence recording allows: neutral mutation overlay and recapitation, both carried out with the aforementioned Python packages rather than SLiM itself.

Neutral mutation overlay adds neutral mutations onto the trees of a tree sequence, sprinkling them randomly according to the neutral mutation rate in each genomic region and the lengths of the branches in each tree. This is exactly equivalent to forward-simulating the neutral mutations, even when the forward simulation has non-neutral dynamics; one can therefore omit the neutral mutations from the forward simulation (as we already did in model 2) and add them afterwards [23,44]. Overlaying neutral mutations after simulation is much more efficient – typically an order of magnitude or more – because they only need to be overlaid onto the branches of the evolutionary trees that lead to the final generation of the simulation. All other branches of the trees went extinct, and were already trimmed away by simplification.

Additionally, all of the overhead of tracking the neutral mutations during forward simulation – carrying them forward into each new generation while shepherding them through the process of recombination – is no longer necessary. We already witnessed the performance benefits of this approach in model 2, since it omits neutral mutations; however, with tree-sequence recording we will regain neutral mutations in the final results of the simulation, while paying very little additional performance cost for doing so (beyond the cost of tree-sequence recording itself).

Recapitation is the process of constructing a neutral burn-in history for a simulation’s initial population using the coalescent, leading backwards in time from the beginning of a forward simulation back to the point of coalescence for each position along a chromosome. It allows you to skip doing a neutral burn-in for a forward simulation; you can start your forward simulation at the end of the burn-in period, without actually simulating the burn-in, and then after your forward simulation has finished you recapitate the burn-in’s neutral history. This is much more efficient than forward-simulating the burn-in of 10*N* generations or more (see above), in part because the coalescent is so fast, but also because recapitation does even less work than a normal coalescent simulation would; it only needs to coalesce backwards from the particular branches that lead to the final generation of the forward simulation. Again, all the other branches that went extinct have already been trimmed away by simplification, which substantially reduces the amount of work needed to simulate the coalescent process [23,30].

However, in this model the burn-in is non-neutral, so substituting a neutral coalescent burn-in for it will be an approximation. The impact of this will depend upon the proximity to the present; sufficiently far back in time, making the burn-in neutral probably has little effect (since little signature of the fine details of the evolutionary process would survive from that far back in time). Closer to the present, the signature left by neutral versus non-neutral dynamics in the pattern of genetic diversity would be larger. One can perhaps therefore draw a line, partway through the burn-in period, and simulate neutral dynamics back in time from that line, and non-neutral dynamics forward in time from that line. Where to draw that line is a difficult question that will depend upon your goals, and is probably best resolved empirically: try drawing the line at different times, and see how close to the present you can draw it without it significantly distorting the patterns of genetic diversity you’re interested in, whatever they might be (the level of diversity, the distribution of lengths of runs of homozygosity, patterns of linkage, and so forth).

The closer to the present you draw the line, the more speed benefit you will derive from using recapitation, but the more likely it is that your results will be distorted by the substitution of neutral dynamics in place of non-neutral dynamics. Many simulations do a neutral burn-in anyway, and so recapitation can substitute for most or all of that burn-in, for a performance gain that is often more than an order of magnitude.

In our model here, we will simply draw the line for recapitation at generation 65795, leaving *N* generations of non-neutral burn-in at the end of the burn-in period, and doing the remaining burn-in (back to coalescence; probably more than 9*N* generations) with recapitation, substituting neutral dynamics for the original non-neutral dynamics of models 1 and 2.

With that preamble, we can look at the code for model 3 in Code Sample 3. Only differences from model 2 are shown, and there are very few.

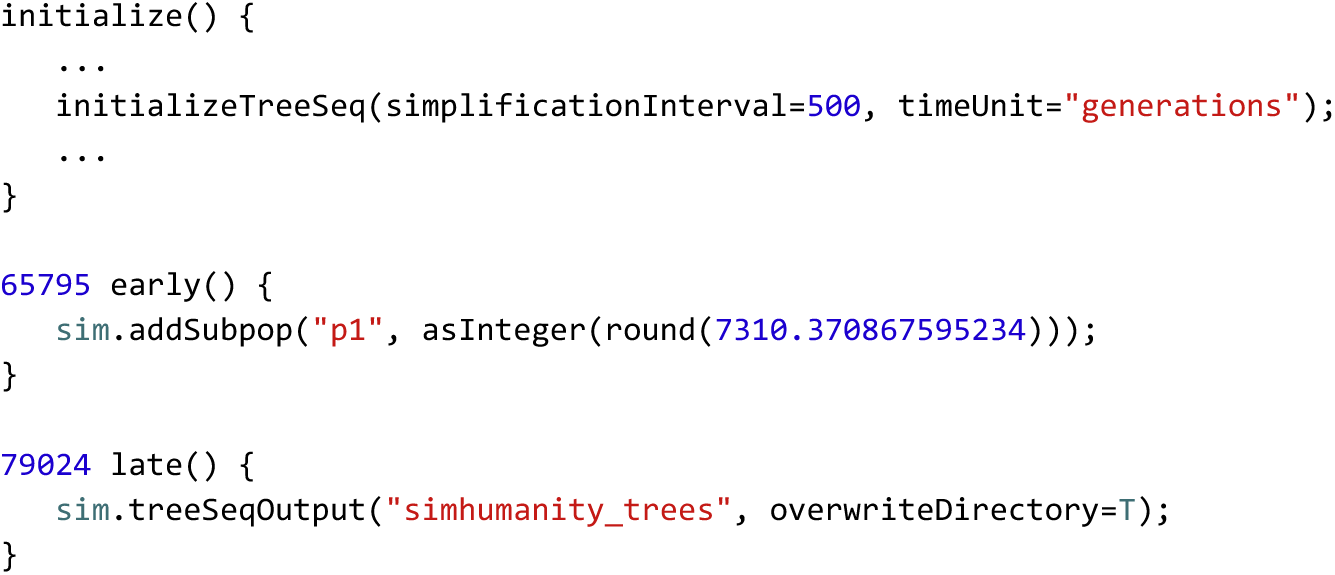

**Code Sample 3.** This version of the SimHumanity model adds tree-sequence recording. Only differences from model 2 are shown here; the full model is provided in `model3_treeseq.slim` in our repository. The `65795 early()` event replaces the `1 early()` event of model 2, starting forward simulation 9*N* generations later than model 2 did; the rest of the burn-in will be conducted with recapitation in Python as discussed in the text. The `79024 late()` event here replaces the (empty) end event in model 2. As before, comments are omitted here, but are included in the model in the repository.

We have added a call to `initializeTreeSeq()` in the `initialize()` callback to turn on tree-sequence recording. The `simplificationInterval` parameter here is optional; we are telling SLiM to simplify every `500` generations. If we didn’t tell it that, it would automatically simplify according to a schedule that it would determine heuristically, which would probably also be fine; performance is not extremely sensitive to the simplification interval, because the time taken by simplification is roughly proportional to the size of the tree sequence; the less often you simplify, the longer each simplification takes, so it tends to balance out for intervals of 250 to 1000; outside of that range, it does often tend to get slower. The `timeUnit` parameter labels the timescale of the tree sequence as being one generation per tick (true because this is a Wright–Fisher model with non-overlapping generations), providing information that improves downstream compatibility with `tskit` and `msprime`.

We have also added a call to `sim.treeSeqOutput()` at the end to write out the trees archive to a directory named `simhumanity_trees`. The `overwriteDirectory` parameter there just tells SLiM that if a trees archive is already present at that path it should be overwritten, unless it appears unsafe to do so (if there are other files present inside the directory besides `.trees` files, for example).

The only other change is that we create the initial `p1` subpopulation, of size `7310` individuals, in generation `65795` instead of generation `1`. This gets rid of the first 9*N* generations of burn-in, which we will do with recapitation instead as discussed above. We could just as easily move the scheduling of all of the other events in the model backwards by 65794 generations; this change is just easier and makes for fewer differences relative to model 2.

A modified version of model 3, found at `model3_treeseq_FIGURE.slim` in our repository, was used to generate the data presented here; its main difference from model 3 is that (like the figure-producing versions of the other models) it sets a random number generator seed of 1, for reproducibility. A run of this version of model 3 at the command line, which produced the `simhumanity_trees` trees archive provided (see Data availability), took 16472 seconds (4.58 hours) and used 78.0 GB of memory, running on the same test machine. At the end of the run there were 620,596 mutations segregating (99.99% of which were deleterious, 0.0135% beneficial) and 85 fixations (29.41% of which were deleterious, 70.59% beneficial). By comparison, model 2 above had 625,065 mutations segregating (99.98% deleterious, 0.0152% beneficial) and 6028 fixations (94.77% deleterious, 5.22% beneficial). From this comparison it can be seen that model 3, due to its curtailed burn-in period, unsurprisingly had far fewer fixations overall; this means that the patterns of genetic diversity at the end of the model run (after recapitation and neutral mutation overlay) might not show the correct signatures of past sweeps and background selection, although that signature should be present for the final 1*N* generations of burn-in, at least. Model 3 did reach something like the equilibrium number of segregating mutations of both types, however – although perhaps not yet at an equilibrium frequency distribution, as we examined with model 1. Whether these possible differences from the full non-neutral burn-in of model 2 mean that this 1*N* non-neutral burn-in period is insufficient would depend on the research questions being addressed by the simulation; the data relevant to the question of interest would need to be examined more closely.

The `simhumanity_trees` archive produced by this model is 27.1 GB. The `.trees` file for chromosome 1, the largest, is 1.61 GB; for chromosome 21, the smallest of the autosomes, it is 0.761 GB. The relatively small difference between these sizes, despite the fact that chromosome 21 is about one-fifth the length of chromosome 1, is likely due to the fact that all of the chromosomes share the same node table (part of the data produced by tree-sequence recording); that node table is very large, and is duplicated in each of the `.trees` files, greatly increasing the overall size of the trees archive. Perhaps as a result of this large-scale data duplication, the trees archive compresses down to 6.89 GB, almost a 75% savings; the compression algorithm perhaps noticed the duplicated node tables. Nevertheless, because of the very large size of this trees archive even after compression, we do not distribute it in our data repository; readers who wish to obtain it can generate it themselves by simulation, or can download it from the separate link provided in “Data availability”.

Having produced or obtained the trees archive, we can now perform recapitation and neutral mutation overlay in Python to complete the burn-in process for the simulation. The code for doing so is shown in Code Sample 4.

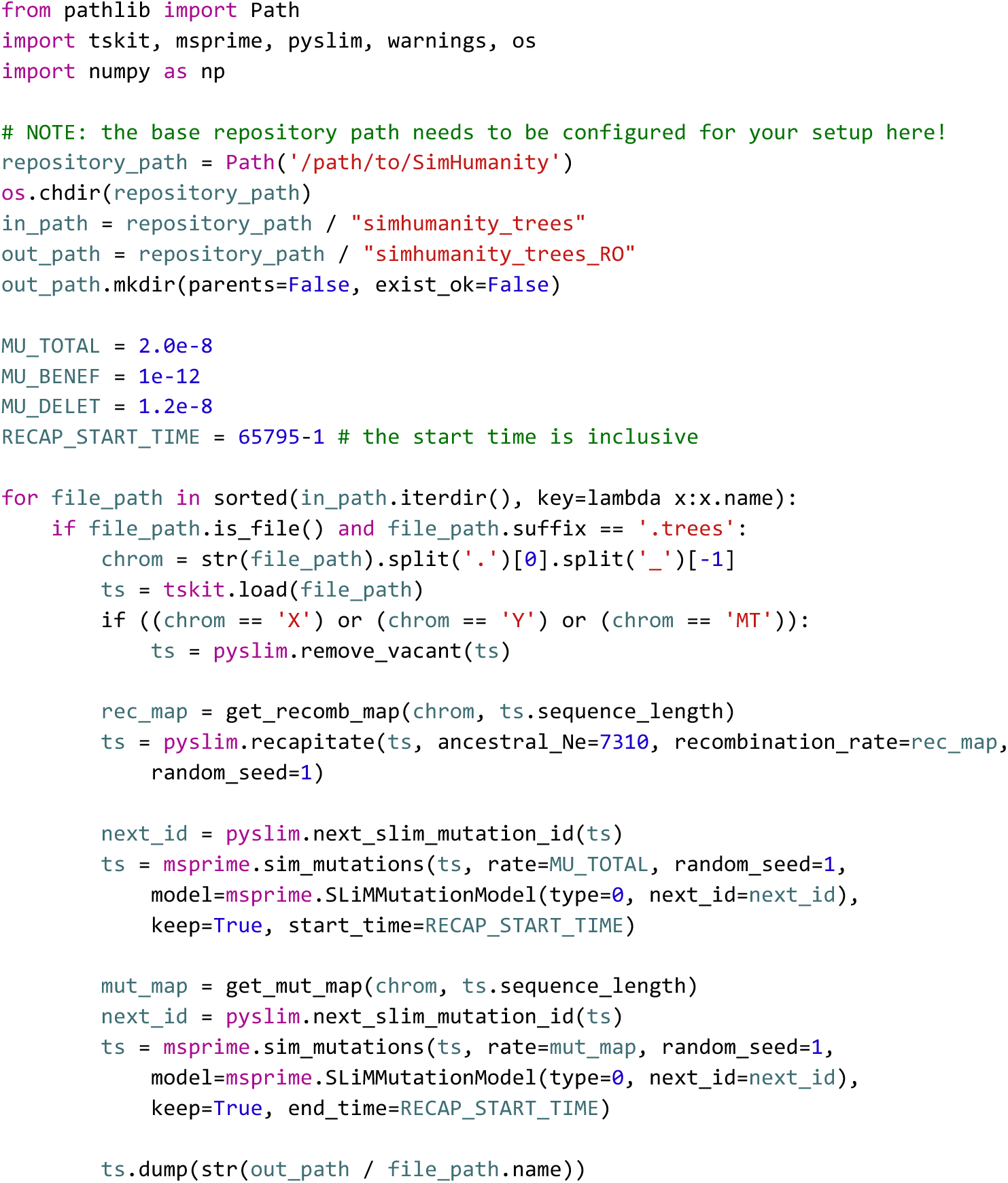

**Code Sample 4.** Python code to perform recapitation and neutral mutation overlay upon the trees archive produced by model 3. This script reads each chromosome’s tree sequence from the `simhumanity_trees` trees archive at the top level of the data repository (note that it needs to be downloaded separately; see “Data availability”) and then recapitates and overlays neutral mutations. The resulting tree sequences are written out to a new trees archive named `simhumanity_trees_RO`. This script is available in our repository as `model3_analysis.py`. As before, elements such as comments, output code, and timing code are omitted here for brevity, including the definitions of the `get_recomb_map()` and `get_mut_map()` functions; all of these elements are included in the full script provided in the repository. A random number generator seed of 1 is used for each of its stochastic steps for reproducibility. See the text for further discussion.

The Python code is fairly straightforward; after importing necessary packages, it loops over the `.trees` files in the trees archive and processes each one. That processing begins by reading the `.trees` file from the trees archive. Next, vacant nodes are removed; these are placeholders for chromosomes that are not modeled in a given individual, such as the Y chromosome in females, and thus only need to be removed when loading the X, Y, and mitochondrial chromosomes where vacant nodes might be present. The recombination rate map for the chromosome is then read in, in much the same way as in the SLiM model, and is used to recapitate the tree sequence to produce the burn-in period’s ancestry back to coalescence.

Neutral mutation overlay is then done in two stages, mirroring the structure of the burn-in period. For the neutral burn-in period, from the time at which recapitation begins back to coalescence, neutral mutations are overlaid using the total mutation rate of the model, `MU_TOTAL`. For the final 1*N* generations of the burn-in, which were forward-simulated in SLiM with deleterious and beneficial mutations (Code Sample 3), neutral mutations are overlaid in such a manner as to supplement what SLiM already simulated: with a rate of `MU_TOTAL` in non-coding regions, and with a rate of `MU_TOTAL - MU_BENEF - MU_DELET` in coding regions where deleterious and beneficial mutations were already simulated. The details of configuring the mutation rate map in this manner are in the `get_mut_map()` function, not shown in Code Sample 4 but supplied in the complete `model3_analysis.py` script provided in the repository. For both of these neutral mutation overlay operations `msprime`’s SLiM mutation model is used to produce new mutations that are compatible with the existing mutations from SLiM, for simplicity.

This script writes out a new trees archive, `simhumanity_trees_RO`, that contains the tree sequences resulting from recapitation and neutral mutation overlay. This trees archive is 36.2 GB (9.57 GB compressed), so these operations added about 9.1 GB of data to the tree sequences. This is mostly in the form of 129,950,887 overlaid neutral mutations, 79,668 of which occur during the recapitated portion of the simulation, the remainder during the SLiM portion. This Python code took 8175 seconds (2.27 hours) to run on the same test machine. The total runtime for the whole simulation pipeline is therefore 6.85 hours. Compared to model 2’s runtime of 19.4 hours, the full model 3 pipeline runs in about 35.3% of the time.

Model 2 used 25.8 GB of memory, whereas model 3 used 77.8 GB; the difference is due to tree-sequence recording, which can use quite a lot of memory. Here we set a simplification interval of 500 generations; a shorter interval would likely decrease the memory usage, although that might (or might not) come at a significant cost to runtime. As mentioned earlier, the optimal balance between runtime and memory usage will depend upon your circumstances; there is no one correct simplification interval.

The benefits of tree-sequence recording might seem small in this instance; model 3 was only three times as fast as model 2, and it used about three times as much memory as model 2. This comparison is misleading, however. At the end of a run of model 2, we have only the non-neutral mutations; but at the end of the model 3 pipeline, we additionally have the neutral mutations, as well as the complete ancestry trees for every location in the genome. Both of these provide valuable additional data; and yet there was also a larger performance advantage than is obvious. As discussed before, a run of model 2 that included neutral mutations took 36.93 hours with a peak memory usage of 48.2 GB – and even then, it still did not provide the ancestry trees that tree-sequence recording provides. The use of tree-sequence recording with recapitation and neutral mutation overlay thus effectively runs in 18.5% of model 2’s time (more than five times faster), with peak memory usage only 1.60 times higher, and provides the tree sequences as additional output, at the price of the approximated burn-in period as discussed above.

After the recapitation and neutral mutation overlay performed by the script in Code Sample 3, the final analysis of the simulation results was done in a separate Python script, `model3_figure.py`, that is provided in the repository but is not shown here (its implementation details are not our focus). The resulting plot is shown in Fig. 3.

**Figure 3.**
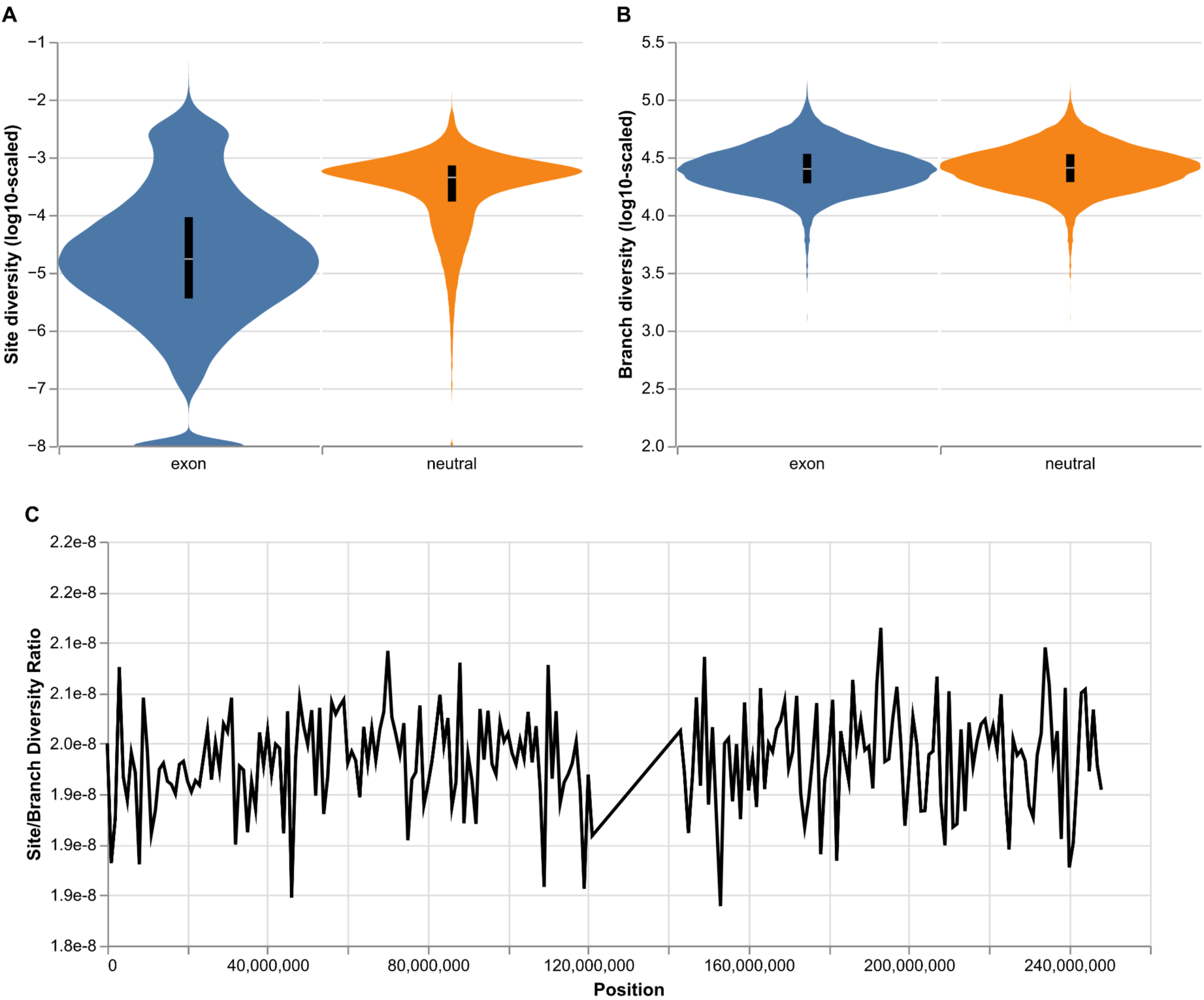
Site- and genealogy-based genetic variation summaries from model 3 (Code Samples 3 and 4). (A) Distribution of site diversity within exons and neutral regions across the autosomes. Each stretch of contiguous sites belonging to the same category (exon or neutral) was considered a single observation. (B) Distribution of branch diversity within exons and neutral regions across the autosomes. (C) Ratio of site to branch diversity statistics for all 1Mb windows along chromosome 1.

With tree sequences, it is possible to compute two different summaries of genetic variation: by site, which is the typical nucleotide diversity (*π;* mean number of pairwise differences per site) computed from genetic data, and by branch, which is the mean pairwise branch length separating samples and equates to the expectation for the site statistic under the infinite sites model of mutation (Ralph et al. 2020) [45]. In Fig. 3A, we show that site diversity is sensitive to selection, such that exonic regions display reduced variation. On the other hand (Fig. 3B), branch diversity is robust to the effect of selection. This result is somewhat surprising, given that both positive and negative selection affect tree shape and reduce tree height. However, the strongly deleterious mutations simulated here were likely removed before having substantial effects on the trees, whereas beneficial mutations are sufficiently rare (as seen in Fig. 2) that their sweep signatures might be lost or averaged out across the many exons where a sweep did not occur. In Fig. 3C, we show that the ratio of site to branch diversity hovers around its theoretical expectation of 2e-8, the total mutation rate. Interestingly, the variation along the chromosome is much smaller than what Ralph et al. (2020) [45] saw in real human genetic data, which may be explained by misspecification of our selection model or mutation rate variation, or by the fact that much of the burn-in of model 3 was done neutrally, with recapitation, rather than forward-simulated with deleterious and beneficial mutations. This analysis thus illustrates the additional power provided by the ancestry information represented in the tree sequences produced by model 3.

## Discussion

We have shown three versions of a model, SimHumanity, that is designed to simulate the complete human genome, including autosomes, sex chromosomes, and mitochondrial DNA.

The first version of the model sets up the genetic structure and then simulates a simple fixed population size of *N*=1000 for 10*N* generations. For this model, we showed how the site frequency spectrum (SFS) changes over time as the simulation builds up towards an equilibrium level of genetic diversity (Fig. 1), illustrating the importance of a “burn-in” period to reach such an equilibrium before beginning non-neutral dynamics.

In the second version of the model we removed neutral mutations from the model for speed, and then added the demography of the Gravel et al. (2011) [25] out-of-Africa model to demonstrate how SimHumanity can be customized to include demographic events. For this model we showed that, given the mutation rate and distribution of mutational fitness effects, selective sweeps often overlap in time (Fig. 2), which could be important to their evolutionary dynamics as a result of Hill–Robertson interference [46–49].

In the third version of the model we added tree-sequence recording, which allowed us to utilize the techniques of recapitation and neutral mutation overlay for greater speed. Recapitation operates after the simulation has finished to construct a neutral burn-in history for the starting population by proceeding backward in time to coalescence, and neutral mutation overlay then adds neutral mutations onto the recorded ancestry trees; both techniques made the simulation much faster than it would have been if the burn-in and neutral mutations had been forward simulated directly in SLiM. However, as discussed for that model, it can be problematic to substitute a neutral burn-in in place of a non-neutral burn-in; it might be best to convert only part of the burn-in period to neutral, or to perform a staged burn-in in some manner, to avoid artifacts due to the optimization techniques being used. Beyond improving performance, we highlight the usefulness of the tree sequence object for downstream analyses, which is enabled by the `tskit` and `pyslim` Python libraries. As a proof of concept, we compare different summaries of genetic variation within coding and non-coding regions (Fig. 3).

As illustrated by the runtimes of these three models, the forward simulations are often quite long – taking hours, days, or even longer (also see [37] and [36]). This can be prohibitive, especially for large genomes or large population sizes, when many replicate runs are needed, or when a high-dimensional parameter space needs to be explored. Tree-sequence recording is one technique that can speed up forward simulations, as shown in model 3, through the techniques of recapitation and neutral mutation overlay as discussed (also see the `pyslim` documentation [50], and [51]).

Several other techniques for achieving greater simulation speed are not demonstrated in our three example models, but warrant brief discussion. The first is model rescaling [30,52–55]. Model rescaling reduces the population size by some rescaling factor *Q*, and rescales other properties of the model like the mutation rate, the recombination rate, selection coefficients, and the number of generations using *Q* in an effort to keep the model in balance, such that it produces similar results to the unscaled model. In some cases this can work quite well, even with a fairly large rescaling factor, greatly reducing simulation times; in other cases even a small rescaling factor, such as *Q*=2, can introduce detectable artifacts that might not be acceptable.

Recent work has begun to characterize these effects of rescaling [30,55], but a great deal remains to be understood about it, and so it must be used with caution; with that caveat, it can be a very powerful technique for achieving better performance.

One area where model rescaling can be particularly effective is during the burn-in period. In model 3, the first part of the burn-in is entirely neutral and done with recapitation, and the last part of it is forward-simulated with both beneficial and deleterious mutations. Such schemes can be more complex, and can include rescaling that is also implemented in a multi-stage fashion. For example, one could use recapitation to provide the simulation’s deep-time ancestry, then switch to forward simulation that includes only deleterious mutations with a large rescaling factor *Q*, then switch to forward simulation that also includes beneficial mutations and uses a smaller *Q*, and then finish off the most recent segment of the burn-in, closest to the present, with no rescaling at all. Neutral mutations could be overlaid across this whole history (with appropriately rescaled mutation rates in the respective burn-in periods, following the different *Q* values) after forward simulation has completed. We are not aware of any research that has focused on the effects of such complex burn-in schemes, yet such techniques seem very promising for bringing large simulations, including simulations of human evolutionary history, within reach.

Another technique to mitigate the long runtimes of forward simulations is the use of a “surrogate model” such as a Gaussian Process that is fitted to a SLiM model and can then be used as a substitute for it [56,57], particularly for the estimation of simple summary statistics. Once trained, such a surrogate model can run many orders of magnitude faster than the original SLiM simulation. For example, a surrogate model could then produce simulated data to feed an Approximate Bayesian Computation pipeline; that approach is notoriously data-hungry, often requiring millions of simulations runs, but producing a dataset of that size might be very fast once a surrogate model has been trained. As with model rescaling, however, this technique only approximates the original SLiM model, and so must be employed with caution to ensure that the surrogate model does a good job of representing the behavior of the original model across the whole parameter space used. Nevertheless, this approach holds great promise for making large-scale forward simulations with many replicate runs more practical.

The SimHumanity model that we have presented is intended to provide a simple baseline human model that showcases the ability of SLiM 5 to simulate full human genomes in the context of various models of human demographic history. This basic model serves as a common starting point for more complex scenarios, and naturally has limitations. Some of these limitations fall into the category of “lack of biological realism”; for example, our model does not include pleiotropy, epistasis, polygenicity, regulatory effects, and so forth, and therefore vastly oversimplifies the genomic architecture of humans. Although we do model mitochondrial DNA, we do not model the many thousands of copies per cell that are transmitted each generation via the germline [58], which might affect the mitochondrial DNA’s evolutionary dynamics. Similarly, we make no attempt to model the selective effects of mutations at known sites such as, say, *FOXP2*; we simply model a given distribution of fitness effects across all coding regions in the genomic map. Whether this lack of biological realism is important depends upon the research questions being asked in a particular study; for some questions such detail would be essential to include, whereas for others it would be unnecessary.

In a similar vein, the Gravel et al. (2011) [25] demographic model we used vastly oversimplifies actual human demographic history (and has since been modified and extended in several studies). In particular, the original model was fitted to empirical data assuming neutrality, whereas we used a non-neutral distribution of fitness effects, so we have mixed apples and oranges for pedagogical purposes. Furthermore, we use the effective population sizes from the Gravel model as census population sizes; in a neutral panmictic Wright–Fisher model that would be defensible, but in a model with deleterious mutations, selective sweeps, and population structure, it probably isn’t. In real human populations, factors like assortative mating and social hierarchy affect the effective population size, too, but those factors are not modeled here.

Furthermore, this demographic model does not include the effects of geographic spatiality within subpopulations, such as isolation by distance; each subpopulation is treated as panmictic.

Taken together, the above considerations imply that our model, “SimHumanity”, is in some sense a toy model, hence its tongue-in-cheek name; it is very far from being a true model of humanity! Nevertheless, it can serve as a common baseline for modeling full human genomes using the new capabilities of SLiM 5, and in that respect it constitutes a worthwhile and important step towards greater realism.

Two other points are worth making regarding the limitations of our SimHumanity model. One is that much of its missing realism could easily be added; SLiM can model many of the things listed above. The practical difficulty in doing so lies not in SLiM, but rather the fact that the necessary knowledge is often lacking (of parameter values, of demographic history, of the fitness effects of specific mutations, of epistatic interactions across the genome, etc.); if we had that knowledge, we could improve the SimHumanity model in many respects. Second, perfect biological realism should not be the aim of modeling anyway [59]; that is a fool’s errand. We should instead aim to make models that include only the biological complexity necessary to answer the specific research questions we are asking in a given study; beyond that, additional realism only makes our task harder, not easier. So while the SimHumanity model is limited in many ways, in some cases that is perhaps not the problem that it initially appears to be.

Given SimHumanity as a step forward towards much-needed greater realism in the modeling of human evolutionary history, what are the next steps? One priority for SLiM will be better intrinsic support for more realistic genetics: multiple phenotypic traits in individuals, polygenicity, quantitative traits, pleiotropy, and so forth. Better support for complex genetic architecture in SLiM would allow simulations to dovetail more closely with empirical results from methods such as GWAS and selection scans. It would also allow SLiM to simulate better null models that could provide an improved basis for inference and for the training of machine learning methods.

Another perennial priority for SLiM is performance. As the runtimes of our example models illustrate, individual-based modeling can be frustratingly slow even with the use of techniques such as tree-sequence recording. We have discussed some additional techniques, such as model rescaling and surrogate models, that can mitigate this problem, but they are approximations that must be used with caution, not panaceas. Fundamentally, faster forward simulations would extend the range of simulations that it is possible to run, allowing important research advances. In situations where a large number of replicate runs is needed, SLiM works well on a computing cluster; each replicate can occupy a separate core, fully utilizing the whole cluster. When only one or a few replicates are needed, however, the fact that SLiM is single-threaded (utilizing only one processing core) can be limiting; it is currently not possible to spread a single SLiM run across multiple cores on a cluster to shorten its runtime. This is another area we intend to focus upon in future work, although parallelizing complex software like SLiM is a large undertaking.

In summary, although there is a long road yet to walk, we offer SimHumanity as an important milestone along that road. We look forward to future work that builds upon the foundation of SimHumanity to construct new simulations of human evolutionary history, enabling new discoveries ranging from a better understanding of our relationships to other hominin groups to a deeper appreciation of humanity’s origins in the tree of life.

### Genetic structure data

To build a human genome model, we need genetic structure data – chromosome lengths, recombination rate maps, and maps of the locations of coding sequences. (It would be nice to have mutation rate maps as well, but those empirical data are not yet available, to our knowledge. When they are, they would be easy to incorporate.) Rather than do the legwork to find that data and get it into a form that SLiM can use, we have relied on the software package `stdpopsim` (version 0.3.0; [36,54,60]). This software can run a wide variety of standardized evolutionary simulations across a range of taxa, using either SLiM or the coalescent simulator `msprime`.

We need to state which data sources within `stdpopsim` we used, and provide appropriate citations not only to `stdpopsim`, but to the original sources from which the data have been drawn. For the genome assembly used, they reference the International Human Genome Sequencing Consortium [61]. For the human recombination rate map, they provide more than two dozen maps (many of which are population-specific); we somewhat arbitrarily chose their `HapMapII_GRCh38` dataset, described as “from the Phase II Hapmap project and based on 3.1 million genotyped SNPs from 270 individuals across four populations (YRI, CEU, CHB and JPT)”. They state that “This version is lifted over to GRCh38 using liftover from the HapMap Phase II map previously lifted over to GRCh37.” They reference this dataset as originating with the International HapMap Consortium [62]. For the genome annotation, they provide two choices; we chose their `ensembl_havana_104_CDS` annotation of coding sequences (CDS), described as “Ensembl Havana CDS annotations on GRCh38” and referencing Hunt et al. (2018) [63] as the original source of the data.

For the distribution of fitness effects, or DFE, they provide six choices; we chose their `PosNeg_R24` DFE, which they describe as “The best-fitting DFE simulated by Rodrigues et al. (2024), from among 57 simulated scenarios with varying amounts of positive and negative selection”, further noting that “the shape of the DFE was not inferred, only the proportion of positive and negative mutations”. For the shape of the DFE, they reference Castellano et al. (2019) [64]; for the proportion of positive and negative mutations, they reference Rodrigues et al. (2024) [37]. This DFE defines an overall mutation rate of 2.0e-8. Within coding regions, a rate of 1e-12 is used for beneficial mutations drawn from an exponential distribution, and a rate of 1.2e-8 is used for deleterious mutations drawn from a gamma distribution. Those rates come out of the overall mutation rate, which is the same across both coding and non-coding regions; in other words, within coding regions ∼60% of all mutations generated by the overall mutation rate of 2.0e-8 are deleterious, ∼40% are neutral, and 0.005% are beneficial. Note that a specific demographic scenario in `stdpopsim` can have its own overall mutation rate, following the mutation rate used in the original paper that introduced that demographic scenario; we did not use such a rate, but instead used `PosNeg_R24`’s overall mutation rate, for two reasons. One reason is that our goal is to build a general-purpose human simulator, not to mirror a specific published demographic scenario; and the other reason is that `PosNeg_R24` was fitted to empirical data using the rate of 2.0e-8, so using a different overall rate would produce a distorted DFE.

Having made these choices, we used `stdpopsim` to generate SLiM scripts that would simulate each of the 24 chromosomes individually (considering the X and Y as different chromosomes, as SLiM does). From these generated SLiM scripts, it was then a simple matter to extract the chromosome lengths, recombination rate maps, and CDS annotation maps. The scripts for doing this are provided as `generate_data.txt` and `extract_data.txt` in our GitHub repository (see Data availability), and the final extracted data used by our SLiM models is provided in the directory named `extracted`. See our repository for further details on this process, all within the directory named `stdpopsim extraction`.

We obtained data for the mitochondrial genome (mtDNA) from NCBI, since the version of `stdpopsim` we used did not fully support mtDNA. A length of 16569 bp, a coding sequence annotation, and a FASTA nucleotide sequence (used as the initial mtDNA sequence for our simulations) were all obtained from NCBI Reference Sequence NC_012920.1 (https://www.ncbi.nlm.nih.gov/nuccore/NC_012920.1). The coding sequence annotation was manually converted from 1-based positions to 0-based positions and converted to the desired file format. The FASTA sequence contained one `N` nucleotide; SLiM requires an unambiguous nucleotide sequence, so we arbitrarily changed this nucleotide to an `A`. The resulting data can all be found in the `mtDNA info` directory in our repository. The mitochondrial genome generally does not recombine, so a recombination rate map was not needed.

For the mutation rate of the mtDNA chromosome, we used an average of two recent estimates from the literature. Rebolledo-Jaramillo et al. [65] give an estimate of 1.30e-8, and Zaidi et al. [66] give an estimate of 1.89e-8. Both of these are in units of per base position per year; assuming a generation time of 25 years, the average of these two estimates is a mutation rate of 3.9875e-07 per base position per generation, which we round off to 4.0e-7 for simplicity.

## Data availability

Scripts, data, and our `stdpopsim`-driven methods for obtaining that data are all open source, provided under the MIT license in GitHub at https://github.com/bhaller/SimHumanity. The `simhumanity_trees.zip` and `simhumanity_trees_RO.zip` trees archives produced by model 3 are available separately at http://benhaller.com/slim/simhumanity_trees.zip (6.89 GB) and http://benhaller.com/slim/simhumanity_trees_RO.zip (9.57 GB) respectively, due to their large size.

## Author contributions

BCH and PWM conceptualized the paper. BCH wrote Code Samples 1–3 with support from CWN, and made Figures 1–2. BCH developed the methodology for the extraction of data from `stdpopsim` that is described in “Genetic structure data”. MFR wrote Code Sample 4 with support from BCH, and did the conceptualization, analysis, and coding that produced Figure 3. BCH and CWN wrote the manuscript with assistance from MFR. BCH, CWN, MFR, and PWM participated equally in reviewing and editing the manuscript. PWM and CWN acquired the funding that supported this work, as acknowledged below.

## Acknowledgements

Thanks to the developers of `stdpopsim` for their excellent software, which provided the core data driving our models, and for their helpful support and advice. BCH and PWM were supported by NIH awards R01HG0124734 and R35GM152242. CWN was supported by the Intramural Research Program of the National Institutes of Health (NIH). The contributions of the NIH author(s) are considered Works of the United States Government. The findings and conclusions presented in this paper are those of the author(s) and do not necessarily reflect the views of the NIH or the U.S. Department of Health and Human Services.

